# BLOC1S1 regulates autolysosomal and exosomal dynamics during CD4⁺ T cell differentiation

**DOI:** 10.64898/2026.05.14.725149

**Authors:** Rahul Sharma, Zulfeqhar A. Syed, Sandeep K. Vishwakarma, Kaiyuan Wu, Kim Han, Anand K. Gupta, Christian A. Combs, Michael N Sack

## Abstract

Although the endolysosome system is central to intracellular recycling, signal transduction, and intercellular communication *via* exocytosis, its role in immunoregulation remains incompletely defined. We recently identified that CD4^+^ T cell-specific depletion of BLOC1S1, a component of multiprotein complexes regulating endolysosomal biology, predisposes toward type 2 (Th2) immunity. We therefore hypothesized that the study of BLOC1S1-deficient CD4^+^ T cells would expand our understanding of endolysosomal dynamics in Th2 function. Here, we demonstrate that CD4^+^ T cell BLOC1S1 deficiency resulted in aberrant lysosomal distribution, accumulation of endosomal vesicles, and increased exocytosis, which collectively correlated with enhanced Th2 immune responses. The phenotype was associated with upregulation of key components of the exocytosis machinery, including RAB11 and VAMP7. Functional inhibition of these vesicle trafficking proteins following siRNA knockdown of RAB11 and VAMP7 significantly attenuated Th2 cytokine secretion in BLOC1S1-deficient CD4^+^ T cells, highlighting their essential role in exosome-mediated cytokine export. Furthermore, exosomes derived from BLOC1S1-deficient CD4^+^ T cells promoted Th2 polarization in recipient cells, indicating a mechanism of intracellular amplification. Together, these findings identify BLOC1S1 as a critical regulator of lysosomal dynamics and exocytic vesicle fusion, thereby linking intracellular trafficking mechanisms to Th2 immune regulation.

## 1. Introduction

The regulation of immune cell function is intimately linked to the fidelity of intracellular vesicular organelle systems, including mitochondria, endosomal, and lysosomal networks.[1, 2] In CD4⁺ T cells, mitochondrial homeostasis and fidelity integrates metabolic status with signaling pathways that govern lineage commitment and effector function.[3] Disruption of mitochondrial integrity can initiate pro-inflammatory signaling *via* the release of mitochondrial damage-associated molecular patterns (mtDAMPs), including mitochondrial DNA (mtDNA), which activate cytosolic pattern recognition receptors (PRRs) to drive cytokine production.[4, 5] The integral role of endolysosomal biology in T cells includes endocytic recycling to regulate T cell receptor (TCR) membrane localization and lysosomal control of mTORC1 signaling for T cell activation.[2] Moreover, exosome-mediated cytokine release contributes to immune regulation, where vesicular trafficking directly influences T cell effector function.[6, 7] Recent data also highlight a role for mitochondria-lysosome contact sites in regulating lysosomal function.[8]

Biogenesis of lysosomal organelle complex 1 subunit 1 (BLOC1S1), an adaptor protein that functions as a component of both BLOC1 and BORC multiprotein complexes, plays a regulatory role in the fidelity and function of intracellular vacuolar organelles.[9–11] BLOC1S1, also known as GCN5L1 or BLOS1, has been most extensively studied in the brain, heart, and liver.[12–16] We have begun to explore its role in T cells to better understand the contribution of vacuolar organelle biology to immune function. Our initial study showed that CD4^+^ T cell-specific BLOC1S1 knockout (TKO) augmented mtDNA leakage into the cytosol, resulting in activation of the cGAS–STING–NF-κB pathway and preferential polarization toward the T helper type 2 (Th2) lineage. Functionally, this translated into heightened susceptibility to allergic inflammation in mouse models of dermatitis and asthma.[5] In that study we also identified increased levels of lysosome-associated membrane protein 1 (LAMP1),[5] a key mediator of endolysosomal biology. We therefore posited that the TKO CD4^+^ T cells reveal additional roles of endolysosomal biology in and provide further insight into the BLOC1S1 KO phenotype of preferential Th2 polarization.

In this context, early endosomes mature into late endosomes, which can function as exocytic organelles capable of releasing extracellular vesicles, including exosomes, that modulate immune responses in an autocrine and paracrine manner.[17–19] Exosome secretion depends on Rab family GTPases (RAB11, RAB27A/B), the exocyst complex (EXOC1), vesicle fusion proteins such as vesicle-associated membrane protein 7 (VAMP7), and tetraspanins including CD81 and CD9.[20–23]

Given that BLOC1S1 deficiency perturbs endolysosome trafficking and homeostasis, we hypothesized that it may also impact the dynamics of exosome biogenesis and release, thereby providing an additional mechanism for Th2 immune responses. In this study, we demonstrate, using transmission electron microscopy, immunofluorescence confocal microscopy, and Nano particle tracking analysis, that TKO cells exhibit accumulation of vesicle-rich endosomes, aberrant lysosomal distribution, and increased exosome secretion. Furthermore, BLOC1S1-deficient CD4⁺ T cells displayed increased LAMP1 accumulation, enhanced exosome production enriched for CD81, CD9, and VAMP7. Moreover, exosomes derived from BLOC1S1-deficient T cells supernatants augmented Th2 cytokine profiles when co-cultured with control CD4^+^ T cells, suggesting a mechanism of intercellular amplification of Th2 responses. Genetic depletion studies further demonstrate that this BLOC1S1 deficient phenotype is dependent on RAB11 and VAMP7. Together these data advance our understanding of the emerging role of endolysosome and exosome biology in CD4^+^ T cell activation.

## 2. Methods

### 2.1 Mice

The NHLBI Animal Care and Use Committee approved all animal studies used in this protocol. The mice were maintained on a 12-h light/dark cycle and housed 3–5 mice per cage with free access to water and a standard chow diet (LabDiet). BLOC1S1 CD4^+^ T cell knockout (TKO) mice were generated by crossing BLOC1S1^flox/flox^ mice with CD4-Cre-recombinase mice, as previously described. All mice were generated on the C56BL/6b background. Experiments were performed using 8–12-week-old C57BL/6^flox/flox^ (control) and CD4^+^ TKO mice (backcrossed > 10 generations).

### 2.2 Mouse CD4^+^ T cell isolation and cytokine assay

All *in vitro* assays were performed using between three and five mice per group. CD4^+^ T cells were negatively selected from the splenocytes using CD4^+^ T cell isolation kit (Miltenyi Biotec) and cultured in RPMI 1640 media supplemented with 25 mM HEPES, 10% FBS, and Penicillin/Streptomycin. CD4^+^ T cells (4 × 10^5^ /well in 96-well plate) were activated with plate-coated αCD3 (5 µg/ml, Biolegend) and αCD28 (10 µg/ml, Biolegend) for 3 days. For Th2 differentiation, CD4^+^ T cells (4 × 10^5^ /well in 96-well plate) were cultured under Th2-polarizing conditions using mouse IL-2, mouse IL-4, and rat anti mouse IFNγ (1:100 dilution, STEMCELL Technologies) plates coated with αCD3 and αCD28 for 3 days. Supernatants were collected, centrifuged to remove cells and debris, and stored at -80 °C. The levels of cytokines, including IFNγ, TNFα, IL-4, IL-5 and IL-13 were measured by ELISA (R&D systems). Results were normalized to cell number using CyQuant cell proliferation assay (Invitrogen) or BCA protein assay (Pierce).

### 2.3 Exosome isolation

CD4^+^ T cells were isolated from the spleens of control and CD4^+^-specific BLOC1S1 knockout mice using magnetic negative selection. Purified CD4^+^ T cells were activated on plates coated with anti-CD3 and anti-CD28 antibodies in complete RPMI medium supplemented with exosome-depleted fetal bovine serum. Cells were cultured for 3 days, and conditioned media were collected for exosome isolation. Cell culture supernatants were centrifuged at 2000 x g for 30 minutes to remove cellular debris and exosomes were isolated using the Exo-Quick-TC (System Biosciences) kit following manufacturer’s guidelines. Exosome size distribution was measured by Nanoparticle Tracking analysis using NanoSight system and expression of exosome markers were characterized by western blotting.

### 2.4 RNA Isolation and Quantitative PCR (qRT-PCR) analysis

Total RNA was extracted using NucleoSpin RNA kit (Macherey-Nagel) and cDNA was synthesized using the SuperScript III First-Strand Synthesis System for RT-PCR (Thermo Fischer Scientific). Quantitative real-time PCR was performed using FastStart Universal SYBR Green Master (Roche) on LightCycler 96 System (Roche). Relative gene expression was determined by normalizing cycle threshold (Ct) values to 18S rRNA using the 2^-ΔΔCt^ method.

### 2.5 Immunoblot analysis

Mouse CD4^+^ T cells were lysed using RIPA buffer supplemented with protease inhibitor cocktail (Roche) and phosphatase inhibitors (Pierce). Lysates were separated by NuPAGE 4-12% Bis-Tris gels (Thermo Fischer Scientific) and transferred to nitrocellulose membranes using the Trans-Blot Turbo Transfer System (Bio-Rad). Membranes were blocked with Odyssey Blocking Buffer (Li-Cor) and incubated with primary antibodies overnight at 4 °C. The secondary antibodies conjugated with IRDye 800 CW or IRDye 680RD (Li-Cor) were then applied for 1 hour at room temperature. Immunoblots were scanned using an Odyssey Clx imaging system (Li-Cor) and band intensity was quantified using ImageJ software (NIH). Primary antibodies for LAMP1, CD81, CD9, CD63, RAB11, VAMP7 and GATA3 were provided by Cell Signaling Technology.

### 2.6 Genetic knockdown experiments

Primary CD4^+^ T cells were transfected with 1.5 µM SMARTpool Accell siRNA targeting VAMP7 or RAB11, or Accell control siRNA, in Accell siRNA delivery medium (Dharmacon). Knockdown cells were activated on αCD3/αCD28 (Biolegend)-coated plates for 3 days.

### 2.7 Immunofluorescence staining and microscopy

CD4^+^ T cells were activated as described above and adhered to poly-L-lysine (Sigma Aldrich)-coated glass slides. Cells were fixed with 4% paraformaldehyde (PFA) for 15 min at room temperature, washed three times with PBS, and blocked with 5% BSA and 0.1% Triton X-100 for 1 hour. Cells were incubated with appropriate primary antibody overnight at 4 °C, washed three times with PBS, and then incubated with Alexa Fluor-conjugated secondary antibody for 60 min at room temperature in the dark. Nuclei were counterstained with 4’,6-diamidino-2-phenylinodole (DAPI). Fluorescent imaging was performed using a Zeiss 880 confocal microscope as previously described [5]. Primary antibodies for CD81, LAMP1, RAB11 and VAMP7 were acquired from Cell Signaling Technology and PKH26 dye from MedChemExpress.

### 2.8 Nanoparticle tracking analysis of exosome concentration and size distribution

Exosome samples were thawed on ice and gently mixed to minimize vesicle aggregation prior to analysis. Samples were diluted in sterile, particle-free PBS to achieve the optimal concentration range for nanoparticle tracking analysis (NTA), ensuring accurate particle detection and tracking. All dilutions were freshly prepared immediately before measurement to limit vesicle degradation and surface adsorption. Particle size distribution and concentration were determined using a NanoSight instrument (NS300, Malvern Panalytical) equipped with a 488 nm laser. Instrument settings, including camera level, detection threshold, shutter speed, and exposure time, were optimized at the start of each acquisition session and maintained consistently across all biological replicates to ensure reproducibility. For each sample, five independent videos of 60 seconds duration were recorded at a controlled temperature (22-25 °C). An automated syringe pump was used to maintain a constant sample flow, thereby minimizing particle drift and ensuring uniform sampling. Video data were processed using NTA software with identical batch-analysis parameters applied across all datasets. Key metrics, including mean, mode, and median particle diameters, as well as total particle concentration, were calculated for each replicate. Recordings exhibiting irregular particle movement, excessive background signal, or signs of aggregation were excluded and repeated. Reproducibility across biological replicates was assessed to confirm consistent exosome isolation efficiency from each source. The final results are presented as particle concentration (particles/mL) and size distribution profiles, reported as mean ± standard deviation (SD) across biological replicates.

### 2.9 Statistical analysis

Graphing and statistical analyses were performed using GraphPad Prism 10. Statistical significance was performed using two-tailed paired or unpaired Student t-tests, or two-way ANOVA with Tukey’s post hoc test for multiple comparisons. The *p* value <0.05 was considered statistically significant. Data are shown as mean ± SEM.

## 3. Results

### 3.1 BLOC1S1 deficiency in CD4^+^ T cells results in the accumulation of late endosomes

To begin to explore whether BLOC1S1 plays a role in CD4^+^ T cell lysosome-endosome biology, we employed transmission electron microscopy (TEM). A striking structural feature was the presence of appreciably larger late endosomes with high vesicular content in TKO cells (Figure 1A-C). The individual vesicles visualized by TEM were measured using ImageJ. The vesicles were approximately 50 nm in diameter, irrespective of the presence or absence of BLOC1S1 (Supplemental Figure 1A). To confirm these findings across a larger number of cells, confocal microcopy was employed. Live cells were incubated with the lipophilic fluorescent dye PKH26 to assess intracellular vesicles content. TKO cells showed a substantially higher level of PKH26 accumulation compared to control cells (Supplemental Figure 1B and C). To validate whether these vesicles were exosomes, cells were labelled with fluorescent antibodies directed against the tetraspanin protein CD81. Here, TKO cells showed substantially higher CD81 content (Figures 1D-E). In parallel, we confirm increased LAMP1 levels in the absence of BLOC1S1 (Supplemental Figure 1D and E).

**Figure 1.**
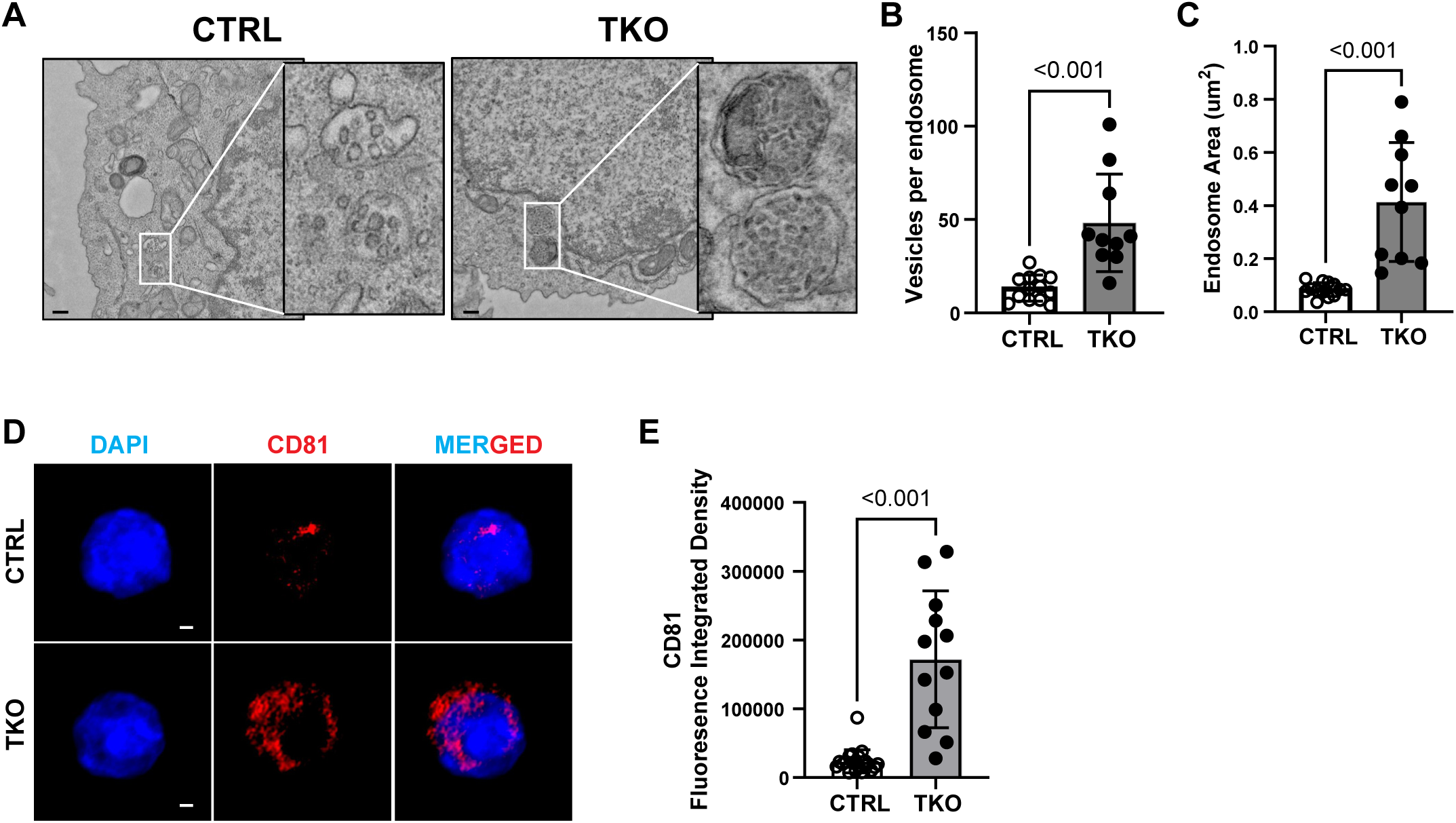
BLOC1S1 deficiency promotes late endosome accumulation and increased CD81-positive vesicular structures in CD4⁺ T cells. **(A)** Representative transmission electron microscopy (TEM) images of control (CTRL) and BLOC1S1-deficient (TKO) CD4⁺ T cells showing late endosomes (boxed regions). Scale bars as indicated (600 nm). **(B)** Quantification of the number of intraluminal vesicles per endosome from TEM images. **(C)** Quantification of endosome area measured from TEM images (µm²). **(D)** Representative confocal microscopy images of CTRL and TKO CD4⁺ T cells stained with DAPI (blue) and anti-CD81 (red). Scale bars, 2 μm. **(E)** Quantification of CD81 fluorescence integrated density. Data are presented as dot plots with mean ± SEM. Each dot represents an individual cell (B-C, E). Statistical significance was determined using an unpaired two-tailed Student’s *t*-test. Experiments were performed independently at least three times.

### 3.2 BLOC1S1 deficiency in CD4^+^ T cells results in increased extracellular exosomes

To determine whether this accumulation of intracellular vesicles was linked to their secretion, extracellular vesicles (EVs) were isolated from the culture supernatants of activated CD4^+^ T cells using a co-precipitation-based method followed by purification, as outlined schematically in Figure 2A. Equal numbers of control and BLOC1S1-deficient CD4^+^ T cells were plated on αCD3/αCD28-coated plates at day 0, cultured under identical conditions for 72 hours, and cell numbers were normalized prior to analysis. The resulting supernatants were used to isolate exosomes which were subjected to nanoparticle tracking analysis (NTA), which provides both particle concentration (particles/mL) and size distribution profiles. Representative size distribution curves and particle concentration from independent replicates are shown in Figure 2B-C. Quantitative analysis revealed a consistent increase (∼ 30-40%) in particle concentration in TKO-derived vesicles compared to controls (Figure 2D-E). In contrast, vesicular size distribution profiles were comparable between groups, with both control and TKO samples exhibiting a predominant vesicle population centered around ∼ 100-150 nm in diameter (Figure 2F), consistent with exosomes. To confirm the identity of these particles, isolated vesicles were analyzed by immunoblot analysis for canonical exosomal markers. TKO-derived vesicles exhibited increased enrichment of CD81, CD9, and CD63, along with LAMP1, compared to control samples (Figure 2G-K), further supporting enhanced exosome secretion in the absence of BLOC1S1.

**Figure 2.**
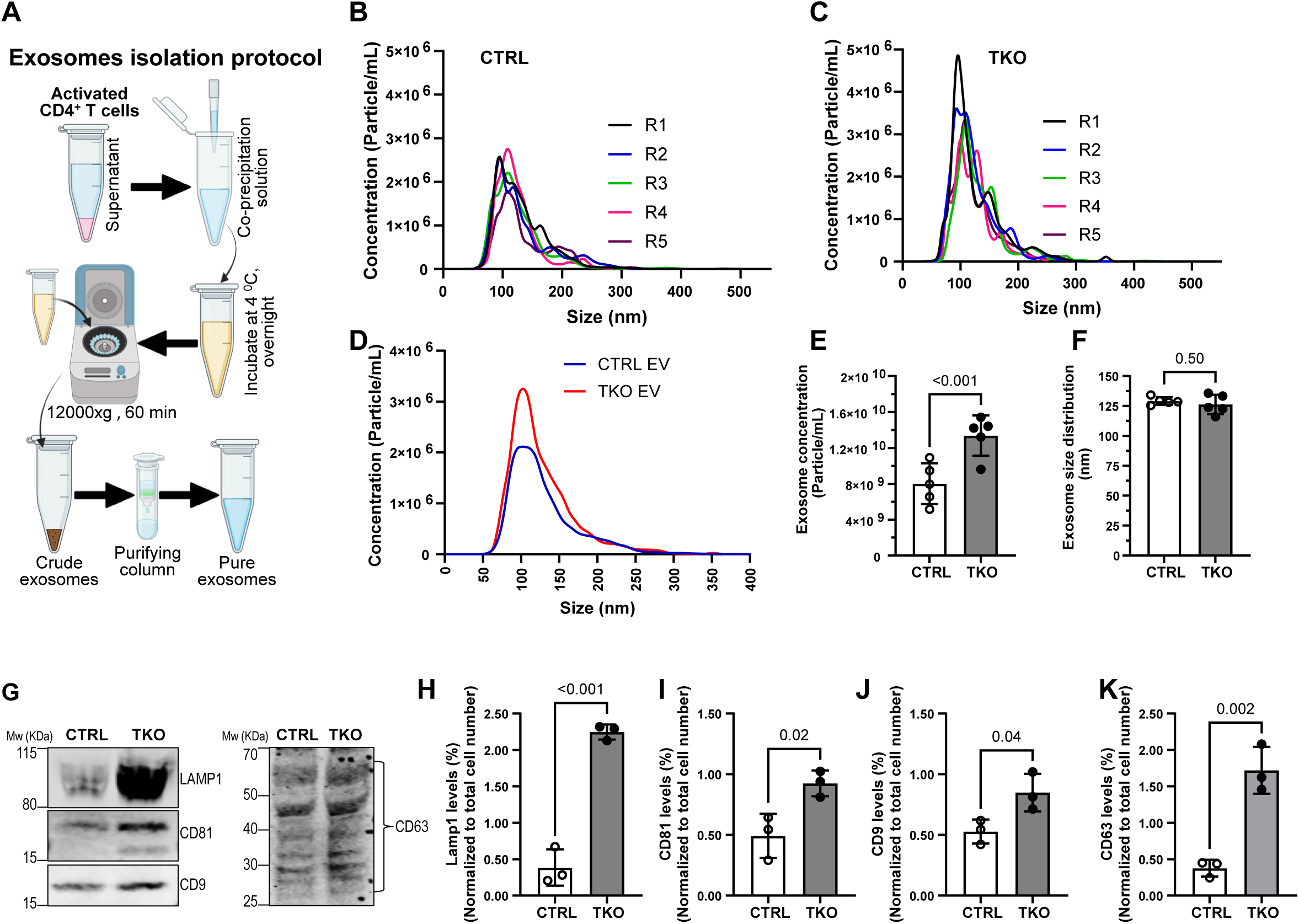
BLOC1S1 deficiency enhances the release of exosome-enriched extracellular vesicles from activated CD4^+^ T cells. **(A)** Schematic overview of the extracellular vesicle (EV) isolation workflow. Activated CD4^+^ T cell culture supernatants were subjected to co-precipitation-based EV isolation followed by purification using a column-based purification method. **(B-C)** Representative nanoparticle tracking analysis (NTA) size distribution profiles of EVs isolated from control CTRL and TKO CD4^+^ T cells from five independent biological replicates. **(D)** Representative overlay of EV size distribution curves from CTRL and TKO samples. **(E)** Quantification of EV concentration measured by NTA (particles/mL). **(F)** Quantification of EV size distribution (nm) between CTRL and TKO samples. **(G)** Immunoblot analysis of EV-associated protein markers from CTRL and TKO CD4^+^ T cell culture supernatants derived EVs. **(H-K)** Densitometric quantification of LAMP1 (H), CD81 (I), CD9 (J), and CD63 (K) protein levels normalized to total input cell number. Data are presented as mean ± SEM with individual data points shown. Statistical significance was determined using an unpaired two-tailed Student’s *t*-test. Experiments were performed with ≥3 independent biological replicates.

### 3.3 Exosomes isolated from BLOCS1-deficient CD4^+^ T cells preferentially drive Th2 polarization

Our prior work showed that the absence of BLOC1S1 in CD4^+^ T cells preferentially promoted Th2 cytokine responses in primary splenic cells and in an ovalbumin-induced asthma model.[5]. To evaluate whether this polarization is recapitulated by TKO-derived exosomes secretion, primary control CD4^+^ T cells were incubated with equal numbers of exosomes from control or TKO CD4^+^ T cells and exposed to TCR activation under Th1- and Th2-polarization conditions. Cytokine analysis showed that exosomes from TKO cells attenuated the production of Th1 cytokines (IFN-γ and TNF-α) and augmented the release of IL-4, a Th2 cytokine (Figure 3A-C). Immunoblot analysis of whole cell lysate from control CD4^+^ T cells treated with exosomes derived from TKO cells showed increased expression of GATA3, a canonical Th2 transcription factor, compared to vehicle or CTRL exosome treated CD4^+^ T cells (Figure 3D-E).

**Figure 3.**
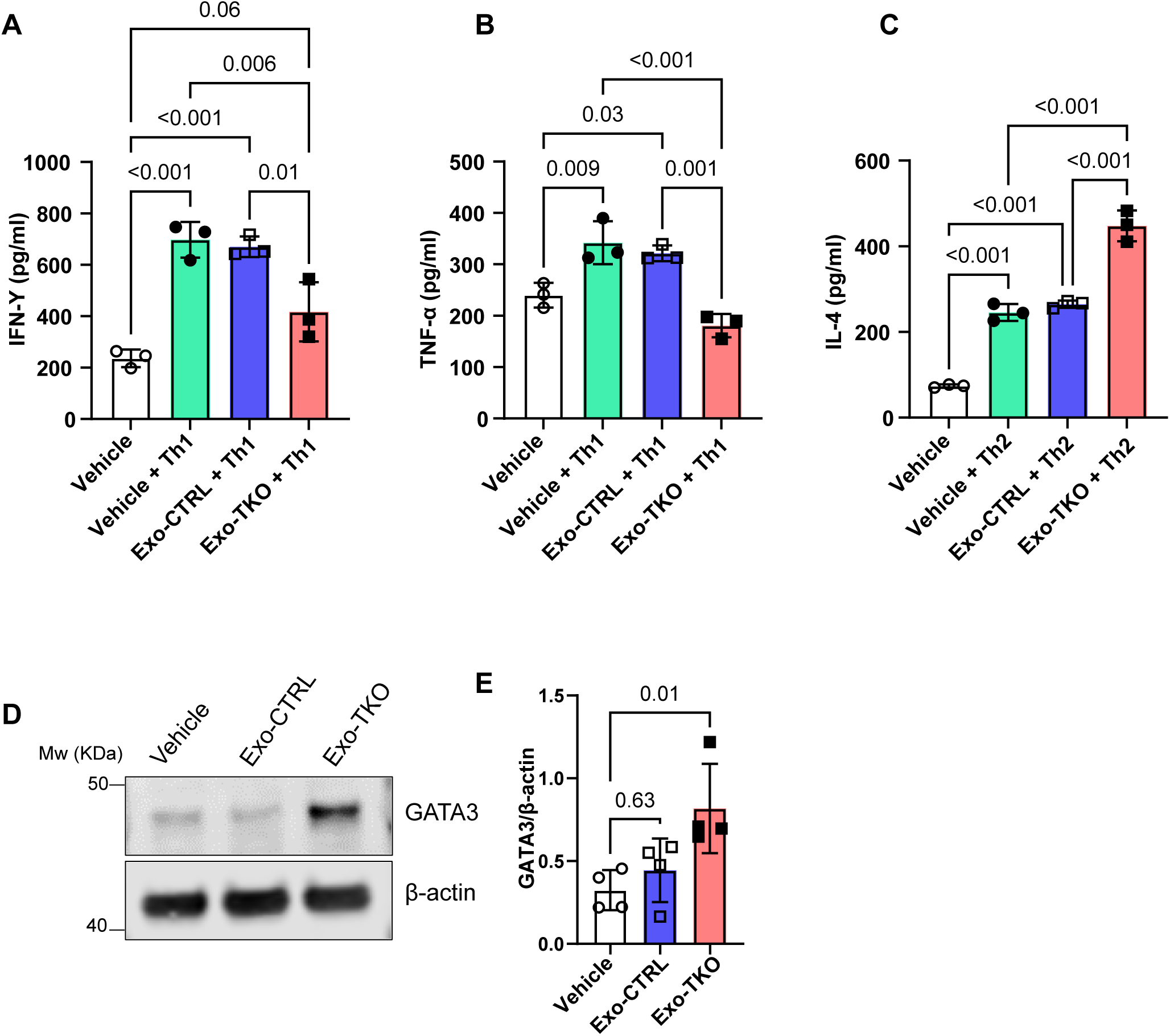
Exosomes derived from BLOC1S1-deficient CD4⁺ T cells preferentially promote Th2-associated responses in recipient CD4^+^ T cells. **(A)** Quantification of IFN-γ production under Th1-polarizing conditions. **(B)** Quantification of TNF-α production under Th1-polarizing conditions. **(C)** Quantification of IL-4 production under Th2-polarizing conditions. **(D)** Immunoblot analysis of whole-cell lysates from recipient CTRL CD4^+^ T cells treated with vehicle, CTRL-derived EVs, or TKO-derived EVs. **(E)** Densitometry quantification of GATA3 protein levels normalized to β-actin. Data are presented as mean ± SEM with individual data points shown. Statistical significance was determined using one-way ANOVA with multiple-comparison testing. Experiments were performed with ≥3 independent biological replicates.

### 3.4 Disruption of RAB11 attenuates Th2 activation in TKO cells

The small GTPase RAB11 functions as a regulator of membrane trafficking, including endosome recycling, vesicular transport to the plasma membrane and exosome excretion pathways.[24] To determine whether RAB11 contributes to the enhanced exocytic and Th2 phenotype observed in BLOC1S1-deficient CD4^+^ T cells, we first assessed its expression in relevant disease contexts. We initially explored public databases and found that gene expression analysis of CD4^+^ T cells from human subjects with asthma (GSE123086 [25]) showed a significant increase in transcript levels of *Rab11* mRNA level compared to healthy controls (Supplemental Figure 4A-C). To validate this *in vivo*, immunofluorescence analysis of lung section from ovalbumin-induced allergic inflammation model demonstrated increased RAB11 signal intensity in TKO mice compared to controls (Figure 4A). At the cellular level, confocal imaging of activated CD4^+^ T cells showed increased RAB11-positive vesicular structures in TKO cells (Figure 4B and Supplemental Figure 4D). This observation was further supported by increased transcript (Figure 4C) and protein levels of RAB11 in TKO CD4^+^ T cells (Figure 4C-D). To determine the functional relevance of RAB11 in T cell activation, RAB11 expression was silenced using siRNA in control and TKO CD4^+^ T cells. Efficient knockdown of RAB11 (Supplemental Figure 4E) resulted in a marked reduction in protein levels of the canonical Th2-associated polarizing transcription factor GATA3 in TKO cells (Supplemental Figure 4F-G). Functionally, RAB11 depletion selectively attenuated Th2 cytokine production, as evidenced by reduced IL-4, IL-5 and IL-13 cytokines levels, while IFN-γ production remained largely unaffected (Figure 4E).

**Figure 4.**
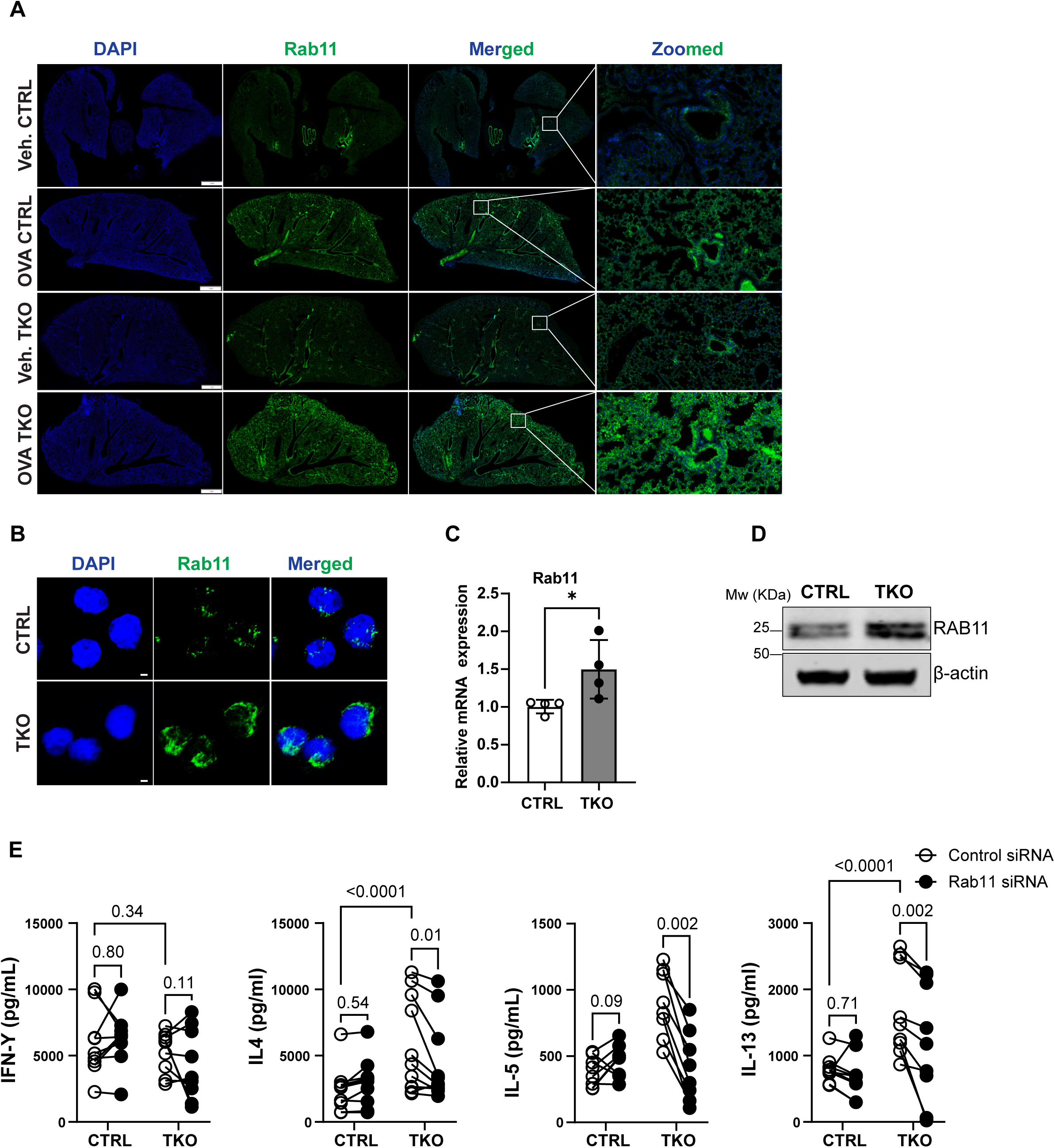
RAB11 expression is increased in BLOC1S1-deficient CD4^+^ T cells and promotes Th2-associated cytokine production. **(A)** Representative immunofluorescence images of lung sections from control and TKO mice (vehicle or OVA-treated) stained for RAB11 (green) and DAPI (blue), with zoomed regions indicated. Scale bar: 100 μm. **(B)** Representative confocal microscopy images of CD4⁺ T cells stained for RAB11 (green) and DAPI (blue). Scale bars, 2 μm. **(C)** Relative Rab11 mRNA expression in CTRL and TKO CD4^+^ T cells by qRT-PCR. **(D)** Immunoblot analysis of RAB11 protein levels in CTRL and TKO CD4^+^ T cells. β-actin was used as a loading control. **(E)** Cytokine production analysis of CTRL and TKO CD4^+^ T cells transfected with control siRNA or Rab11 siRNA. Data are presented as mean ± SEM with individual data points shown. Statistical significance was determined using two-way ANOVA with multiple comparison testing or unpaired Student’s *t*-test where appropriate. Experiments were performed with ≥3 independent biological replicates.

### 3.5 Inhibition of VAMP7 selectively blunts Th2 activation in TKO cells

VAMP7 is a vesicle associated SNARE (v-SNARE) protein that mediates membrane fusion of late endosomal and lysosomal vesicles with the plasma membrane, thereby facilitating vesicle-dependent exocytosis in immune cells.[26] To determine whether VAMP7 contributes to the enhanced secretory and Th2 phenotype observed in BLOC1S1-deficient CD4^+^ T cells, we first assessed its expression and subcellular localization. Analysis of transcript levels revealed a significant increase in *Vamp7* mRNA expression in TKO cells compared to control (Figure 5A). Consistent with this, immunoblot analysis demonstrated elevated VAMP7 protein levels in TKO cells (Figure 5B). To further examine VAMP7 distribution, confocal microscopy was performed. TKO cells displayed increased VAMP7-positive vesicular structures, with prominent accumulation near the plasma membrane compared to control cells (Figure 5C-D). Higher magnification images highlighted peripheral localization of VAMP7-enriched vesicles, consistent with an increased pool of fusion-competent vesicles poised for exocytosis. Concurrently, to determine the functional relevance of VAMP7 in T cell activation, VAMP7 expression was silenced using siRNA in both control and TKO cells. Efficient knockdown of VAMP7 (Supplemental Figure 5A) resulted in a marked reduction in GATA3 in both control and TKO cells (Supplemental Figure 5B-C). Functionally, VAMP7 depletion selectively attenuated Th2 cytokine secretion. Specifically, IL-4, IL-5 and IL-13 production were significantly reduced in BLOC1S1-deficient CD4^+^ T cells, whereas IFN-γ levels were less affected (Figure 5E). These findings indicate that VAMP7-mediated vesicle fusion is a critical regulator of Th2 effector function and supports a model in which enhanced exocytic machinery drives increased cytokine output in BLOC1S1-deficient CD4^+^ T cells.

**Figure 5.**
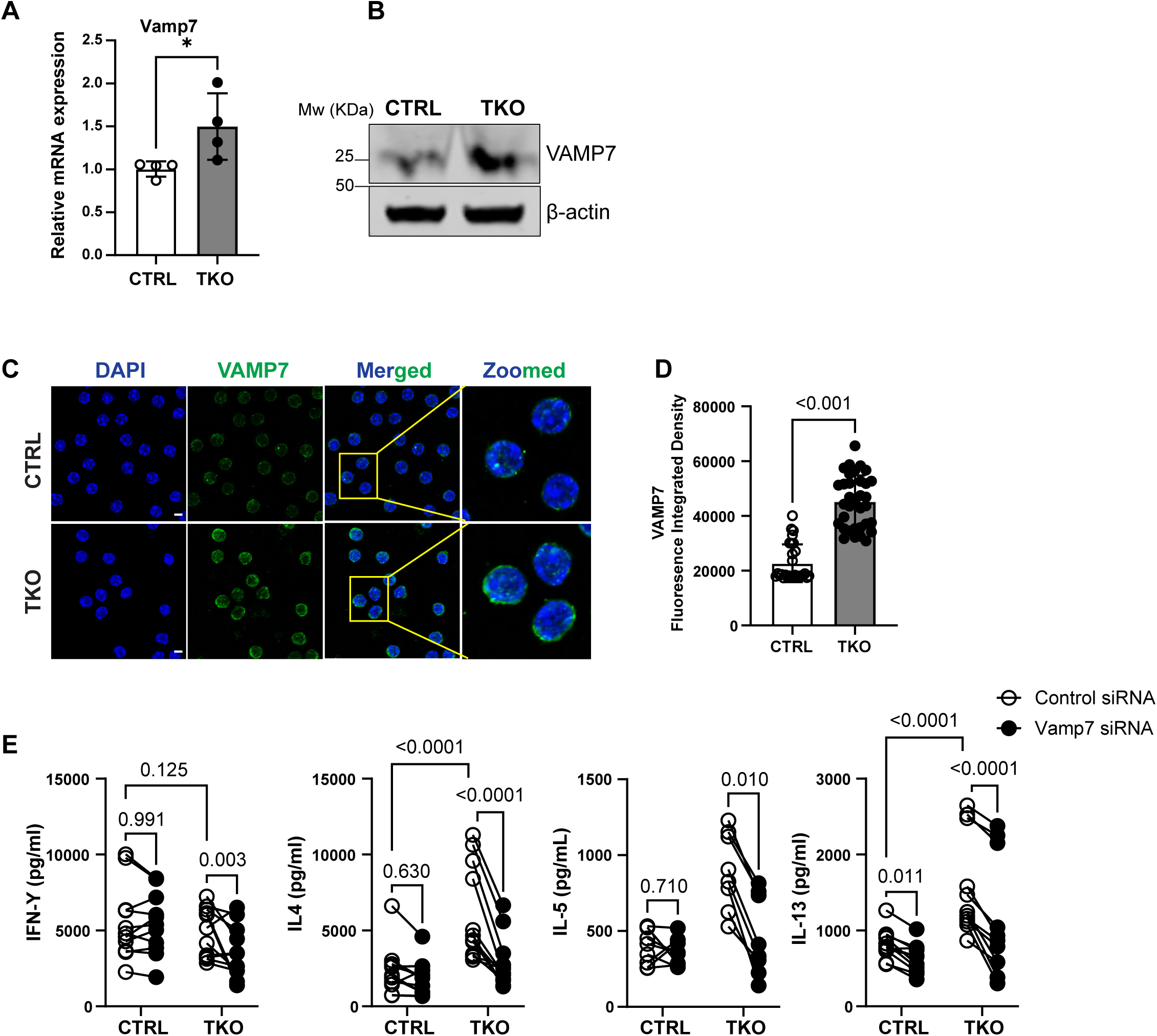
VAMP7 expression is increased in BLOC1S1-deficient CD4+ T cells and promotes Th2-associated cytokine production. **(A)** Relative Vamp7 mRNA expression in CTRL and TKO CD4^+^ T cells by qRT-PCR. **(B)** Immunoblot analysis of VAMP7 protein levels in CTRL and TKO CD4^+^ T cells. β-actin was used as a loading control. **(C)** Representative confocal microscopy images of CTRL and TKO CD4⁺ T cells stained for VAMP7 (green) and DAPI (blue), with zoomed regions indicated. Scale bars, 5 μm. **(D)** Quantification of VAMP7 fluorescence integrated density. **(E)** Cytokine production analysis of CTRL and TKO CD4^+^ T cells transfected with control siRNA or VAMP7 siRNA. Data are presented as mean ± SEM with individual data points shown. Statistical significance was determined using two-way ANOVA with multiple comparison testing or unpaired Student’s *t*-test where appropriate. Experiments were performed with ≥3 independent biological replicates.

## 4. Discussion

In this study we demonstrate that the absence of BLOC1S1 in CD4^+^ T cells results in the accumulation of vesicles within late endosome accompanied by excess exosome secretion. Additionally, incubation of isolated exosomes from BLOC1S1-deficient T cells recapitulates the TKO phenotype by promoting Th2 polarization and cytokine production in control primary CD4^+^ T cells. This study also demonstrated that canonical RAB11- and VAMP7-dependent exocytosis is required to orchestrate this BLOC1S1-deficiency dependent regulation. Together these findings expand our understanding of the contribution of endosome–exosomal dynamics in Th2 polarization.

BLOC1S1 is an adaptor protein that appears as a common subunit in different multiprotein complexes involved in the trafficking and homeostasis of intracellular organelles, including endolysosomes and mitochondria. Structurally, emerging data suggest that two distinct multiprotein complexes, i.e. BLOC-1 and BORC, both incorporate BLOC1S1, may in fact be dynamic with distinct modular assemblies to create structural specialization to meet their diverse functional roles.[27] This structural complexity may contribute to the different phenotypes observed following genetic depletion of individual subunits. In parallel, studies of BLOC-1 and BORC components have linked these complexes to distinct cellular functions, with BLOC-1 primarily associated with early endosome biology and BORC in late endosome biology.[11] BLOC1S1 itself, which is a component of both complexes, has been shown to regulate multiple aspects of vesicular organelle biology, including endosome positioning, synaptic vesicle and lysosome positioning,[28–30] vesicular trafficking into endosomes,[12] lysosomal reformation and lipolysis,[15, 16] and endosomal maturation and function.[12, 31] As the ablation of BLOC1S1 in CD4^+^ T cells differentially promotes type 2 immunity,[5] further study of BLOC1S1-deficient CD4^+^ T cells may provide additional insight into how vesicular programs regulate immune function.

Although understanding of the role of the vacuolar lysosomal-endosome-exosome axis in CD4^+^ T cell biology continues to expand, it remains incompletely characterized. A substantial body of work shows that various forms of endocytosis are involved in αβ[32] and γδ[33] T cell receptor and IL-2Rβ complex[34] recycling, function, and fates. Lysosomal biology similarly regulates mTORC1 signaling,[35] which is instrumental for T cell activation, and recent evidence shows that mis-localization of mTORC1 to late endosomes impairs lysosomal function and contributes to age-associated immune dysfunction.[36] Furthermore, the integrity of endolysosomal homeostasis is required for regulatory T cell suppressive function.[37] The role of exosomes in this system is beginning to be explored, where exocytosis has been shown to contribute to immune synapse function and T cell communication with antigen-presenting cells,[6] and to influence how CD4^+^ T cells modulate CD8^+^ T cells tumor responsiveness.[38] This study advances our understanding of the role of BLOC1S1-deficient exosomes on type 2 immunity.

The study was initiated at the structural level by examining control and TKO cells using transmission electron microcopy and confocal microscopy, while extracellular vesicle content was assessed using nanoparticle tracking analysis. These studies showed that, in the absence of BLOC1S1, vesicles accumulate within late endosomes in parallel with increased extracellular exosome release. Nanoparticle tracking analysis demonstrated that BLOC1S1 deficiency did not affect exosome size. However, incubation with equal numbers of exosomes showed that TKO-derived exosomes augmented h2 polarization in control primary CD4^+^ T cells. Together, these data indicate that loss of BLOC1S1 results in increased late endosomal vesicular accumulation, enhanced exosome secretion, and a greater type 2 immune response. Furthermore, these data suggest that BLOC1S1 regulates late endosome and exocytosis pathways at multiple levels. Future studies will be required to define the molecular content of these exosomes to determine how these exosomes preferentially drive the Th2 effector cell fate.

The biology of exocytosis is complex and dynamic and can be bidirectional through coupling with endocytosis to retrieve extracellular vesicles.[39] Shared molecular components regulate vesicular scission, cytoskeletal trafficking mediators, and membrane fusion. RAB11 acts as a master regulator of trafficking from recycling endosomes to the plasma membrane, whereas VAMP7, a v-SNARE protein, directly mediates fusion of lysosomal and vesicular membranes with the plasma membrane to facilitate exosome secretion. In this study, both RAB11 and VAMP7 were upregulated in the absence of BLOC1S1, and their genetic depletion attenuated canonical signaling pathways and Th2 cytokine production in TKO cells. These findings further support that BLOC1S1 contributes to exocytosis in CD4^+^ T cells and that this pathway is critical in driving Th2 polarization.

The limitations of this study include the need for a more detailed understanding of how BLOC1S1 functions within distinct endolysosomal regulatory complexes. Additionally, the content of these exosomes will need to be defined to determine whether the vesicular modulation of Th2 polarization is driven by RNA, protein, or metabolite cargo.

In conclusion, disruption of endolysosomal homeostasis in the absence of BLOC1S1 results in accumulation of vesicles within late endosomes and increased exosome secretion. These in turn, appears to play an important role in CD4^+^ T cell fate, promoting preferential polarization toward the Th2 lineage. Further dissection of this biology may identify extracellular targets to modulate Th2-driven allergic conditions.

## Acknowledgements

We thank and acknowledge the assistance of the NHLBI Laboratory Animal Core Facility. This research was supported by the Intramural Research Program of the National Institutes of Health (NIH). The contributions of the NIH author(s) were made as part of their official duties as NIH federal employees, are in compliance with agency policy requirements, and are considered Works of the United States Government. However, the findings and conclusions presented in this paper are those of the author(s) and do not necessarily reflect the views of the NIH or the US Department of Health and Human Services. Funding for this study was supported by the NIH intramural programs to MNS (grant HL-005102).

## Conflicts of Interests

All authors declare no conflicts of interest.

## Author Contributions

All authors reviewed the data and contributed to its interpretation, edited the manuscript, and approved the final submitted version. Specific contributions: R.S. conceived, designed, and performed most experiments in the study and drafted the manuscript. Z.A.S. contributed to TEM image acquisition and interpretation and drafting of the manuscript. V.S.K. contributed to data acquisition, analysis and interpretation and drafting of the manuscript. K.W. contributed to data interpretation and drafting of the manuscript. K.H. contributed to data acquisition, analysis, and interpretation and drafting of the manuscript. A.K.G. contributed to data acquisition and interpretation of the data. C.A.C. designed and analyzed light microscopy data and contributed to drafting the manuscript. M.N.S. conceived and designed the study, wrote study protocol and obtained ethical approval, secured funding for the study, analyzed data, and drafted the manuscript.

## Funding

This research was supported by the NHLBI Division of Intramural Research (MNS-ZIA-HL005199).

## Supplementary Figure Legends

**Supplementary Figure 1. Increased PKH26 and LAMP1 accumulation in BLOC1S1 deficient CD4^+^ T cells.**

**(A)** Quantification of vesicle size distribution measured from TEM images. **(B)** Representative confocal microscopy images of CD4⁺ T cells stained with PKH26 (red) and DAPI (blue). Scale bars, 2 μm. **(C)** Quantification of PKH26 fluorescence integrated density. **(D)** Representative confocal microscopy images of CD4⁺ T cells stained with anti-LAMP1 antibody (red) and DAPI (blue). Scale bars, 5 μm. **(E)** Quantification of LAMP1 fluorescence integrated density. Data are presented as dot plots with mean ± SEM. Statistical analysis was performed using an unpaired two-tailed Student’s *t*-test.

**Supplementary Figure 4. Increased RAB11 expression in asthma and functional effects of RAB11 depletion in CD4^+^ T cells**.

**(A-C)** Analysis of publicly available transcriptomic datasets (GSE123086) showing increased RAB11 expression in CD4^+^ T cells from female asthma patients (A), male asthma patients (B), and combined cohorts (C) compared with healthy controls. **(D)** Quantification of RAB11 fluorescence integrated density from confocal microscopy images of CTRL and TKO CD4^+^ T cells. **(E)** Quantification of Rab11 mRNA expression following siRNA-mediated knockdown in CTRL and TKO CD4^+^ T cells confirming efficient RAB11 depletion. **(F)** Immunoblot analysis of GATA3 levels in CTRL and TKO CD4^+^ T cells following transfection with control or Rab11 siRNA. β-actin was used as a loading control. **(G)** Densitometric quantification of GATA3 protein levels normalized to β-actin. Data are presented as mean ± SEM. Statistical analysis was performed using an unpaired two-tailed Student’s *t*-test.

**Supplementary Figure 5. Efficient VAMP7 depletion reduces GATA3 expression in CD4^+^ T cells.**

**(A)** Quantification of Vamp7 mRNA expression following siRNA-mediated knockdown in CTRL and TKO CD4^+^ T cells. **(B)** Immunoblot analysis of GATA3 expression in CTRL and TKO CD4^+^ T cells transfected with control or Vamp7 siRNA. β-actin was used as a loading control. **(C)** Densitometric quantification of GATA3 protein levels normalized to β-actin in CTRL and TKO CD4^+^ T cells. Data are presented as mean ± SEM. Statistical analysis was performed using an unpaired two-tailed Student’s *t*-test.

## Notes

### Competing Interest Statement

The authors have declared no competing interest.

