## Supplementary figures and images for "BLOC1S1 regulates autolysosomal and exosomal dynamics during CD4⁺ T cell differentiation"

### Supp Fig 4

**Supplementary Figure 4**

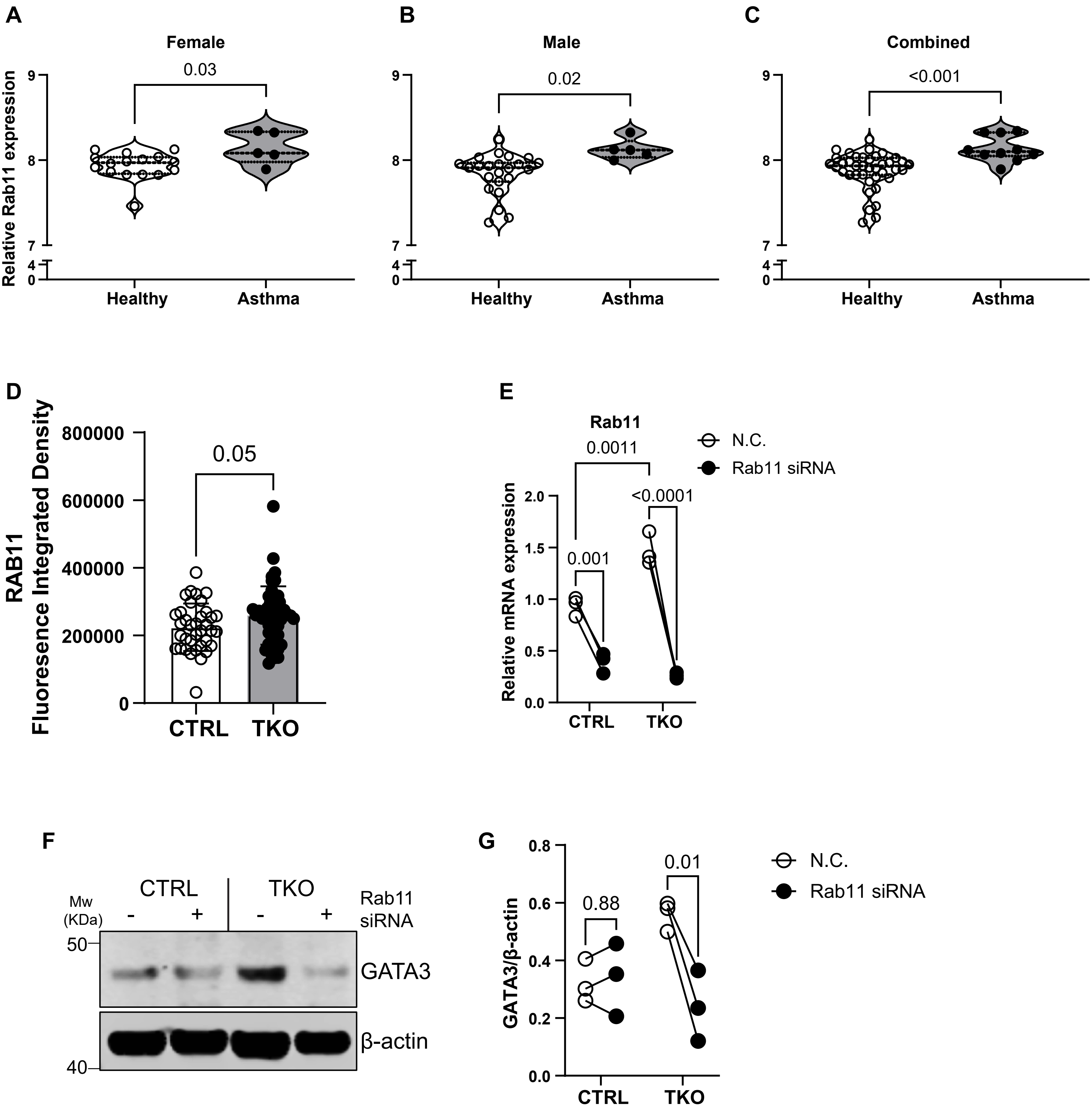

### Supp Fig 5

**Supplementary Figure 5**

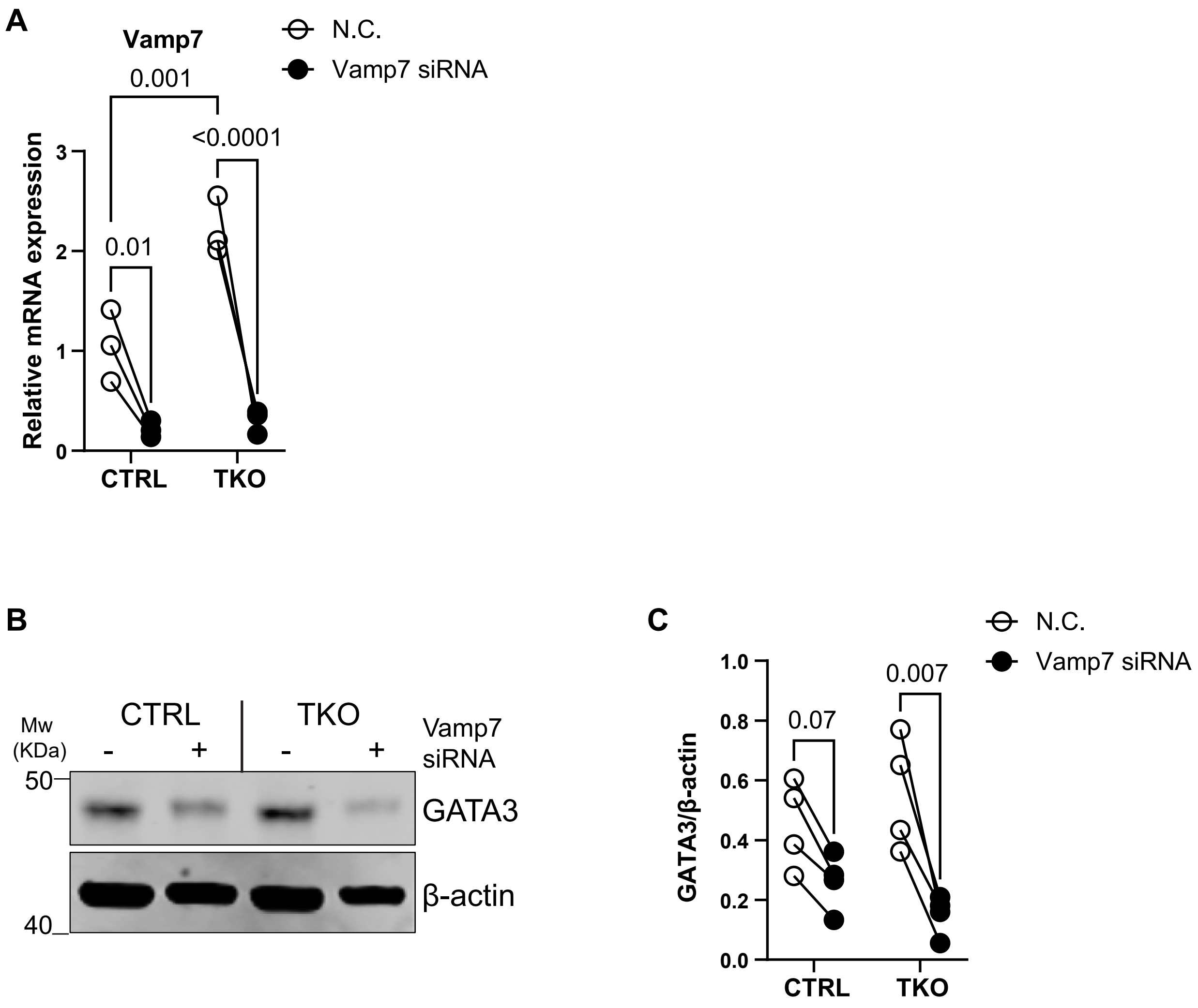

### Supp Figure 1

**Supplementary Figure 1**

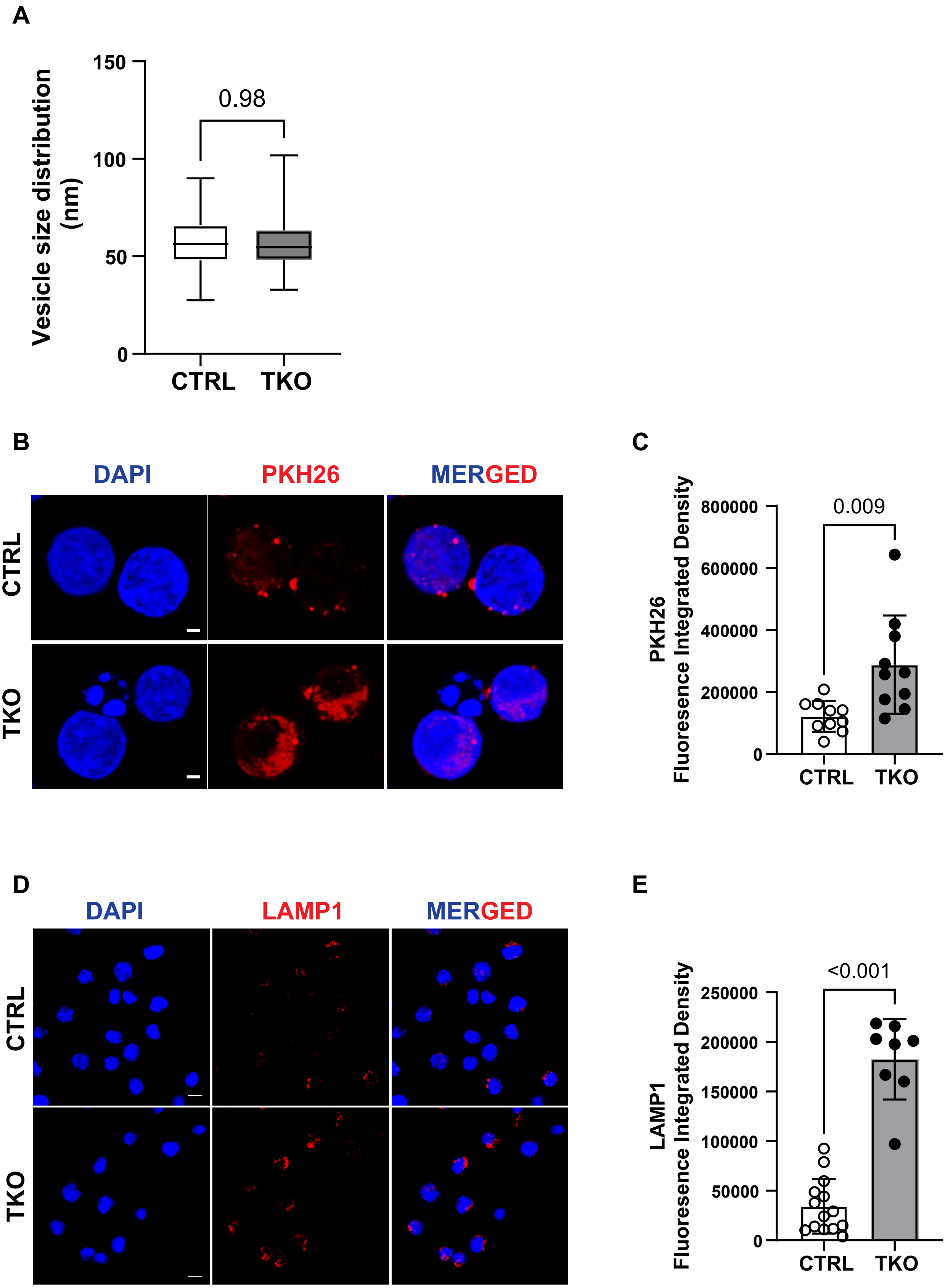
